# Structural brain networks shape individual-level progression of brain atrophy after stroke

**DOI:** 10.64898/2026.05.08.723677

**Authors:** Jing Yang, Yixin Gao, Liyuan Yang, Yaya Jiang, Bin Wan, Şeyma Bayrak, Xinyu Liang, Sofie L. Valk, Maurizio Corbetta, Gaolang Gong

## Abstract

Stroke starts as a focal vascular lesion, but its structural consequences often extend beyond the lesion site, resulting in distributed brain atrophy whose organizing principles remain unclear. Here, using longitudinal MRI data from two stroke cohorts spanning the hyperacute to chronic stages, we tested whether and how post-stroke grey matter volume (GMV) change over time is constrained by large-scale brain networks across individuals. Coordinated deformation modelling showed that longitudinal GMV atrophy patterns were more strongly shaped by structural than by functional connectivity, with structural constraints already detectable in the acute phase. Network diffusion modelling further showed that atrophy patterns at 3 and 12 months were consistent with lesion-initiated propagation on the structural connectome, whereas model performance was weak in the early acute stage. The model-derived propagation stage did not increase monotonically with chronological time, indicating that it reflects the degree to which observed atrophy conforms to a network-consistent diffusion-like pattern rather than elapsed biological time. Inter-individual variation in propagation stage was associated with lesion size and with the topological and molecular properties of lesioned regions. Finally, lesion information combined with network diffusion modelling supported individualized prediction of later GMV atrophy patterns. These findings provide a network-based account of secondary degeneration after stroke and may inform individualized characterization of post-stroke structural reorganization.

## Introduction

Although stroke is initially a focal vascular event, the resulting structural alterations often permeate regions far distal to the lesion site.^1–3^ Neuroimaging studies have shown that grey matter loss, cortical thinning, and white matter abnormalities can emerge in regions remote from the primary lesion, indicating that post-stroke degeneration is not confined to locally injured tissue. These remote changes are clinically consequential, as their extent and spatial distribution have been linked to motor recovery, cognitive performance, and overall functional outcome.^4–8^ A central question, therefore, is how a focal lesion gives rise to distributed brain atrophy over time. Addressing this question is clinically important, because secondary degeneration beyond the lesion may shape the trajectory of recovery, long-term impairment, and the window for targeted intervention after stroke.

Post-stroke atrophy is likely multifactorial, involving not only local ischemic and vascular factors but also inflammatory and metabolic cascades.^9–11^ Beyond these influences, focal injury may also induce downstream abnormalities in regions anatomically connected to the lesion site. ^12–15^ Consistent with this view, remote post-stroke atrophy has repeatedly been observed in regions directly connected to the primary lesion, in keeping with the classical concepts of diaschisis and transneuronal degeneration.^14,16–20^ However, direct lesion-linked connections alone do not fully account for the broader and more distributed atrophy observed across the brain. Accounting for such whole-brain patterns requires considering connectivity not simply as isolated lesion-to-region links, but as a large-scale network that embeds all regions within an interconnected system.

Mechanistic network models, including the coordinated deformation model (CDM) and network diffusion model (NDM), provide a formal framework for testing whether and how large-scale brain networks constrain the spatial distribution and temporal evolution of structural abnormalities across the brain.^21,22^ In neurodegenerative and psychiatric disorders, these models have shown that grey matter abnormalities often follow network-constrained patterns, in some cases with specific regions identified as putative epicentres.^23–26^ However, such studies have so far relied largely on cross-sectional, group-level analyses,^27,28^ providing limited insight into how network-constrained structural change unfolds over time within individuals. In particular, whether whole-brain post-stroke atrophy is similarly constrained by large-scale network architecture remains untested within this modelling framework.

Longitudinal stroke imaging, particularly when combined with mechanistic network modelling, offers a direct opportunity to address this question at the individual level. Stroke also provides a stringent test of the network hypothesis, because lesion location, size, and laterality vary substantially across patients, creating marked heterogeneity in the anatomical starting points of post-stroke degeneration.^16,29,30^ If large-scale network architecture constrains the evolution of post-stroke atrophy, this principle should remain detectable despite such heterogeneity. At the same time, this heterogeneity provides an opportunity to examine how lesion-related factors contribute to inter-individual variation in network-model measures.

In this study, leveraging high-quality longitudinal MRI data spanning multiple post-stroke stages in two cohorts of patients with heterogeneous focal lesions, we first used CDM to test whether longitudinal atrophy patterns were more strongly constrained by structural than by functional connectivity. We then applied NDM to determine whether lesion-initiated network propagation could capture patient-specific atrophy trajectories over time and to characterize inter-individual variation in inferred progression. Finally, we examined how lesion-related factors contribute to this variation and evaluated whether lesion information, together with NDM, could support individualized prediction of post-stroke atrophy.

## Materials and methods

### Participants and datasets

We analysed a longitudinal stroke cohort comprising 132 patients (age range, 19–83 years) with first-ever symptomatic infarction. Participants were not selected on the basis of lesion location, vascular territory, or behavioural phenotype, thereby preserving broad clinical representativeness.^31^ Structural MRI was acquired at approximately 2 weeks (acute baseline), 3 months (subacute stage), and 12 months (chronic stage) post-stroke.

Using previously reported quality-control procedures,^31^ we excluded eight participants because of suboptimal T1-weighted image quality. Patients with isolated infratentorial lesions were additionally excluded, restricting analyses to cortical and subcortical grey matter change. The final analytical sample comprised 72 patients with high-quality baseline structural imaging and at least one follow-up structural scan (72 at baseline, 69 at 3 months, 62 at 12 months). Demographic and clinical characteristics are summarised in Supplementary Table 1.

We also analysed an independent early-phase cohort (n = 28) from a previously published dataset,^32^ which provided coverage of the first week after stroke onset and complemented the main longitudinal cohort beginning at approximately 2 weeks post-stroke. MRI was acquired at Day 0 (<24 h), Day 1 (24–48 h), and Day 5 (4–6 days; mean ± SD = 4.93 ± 0.38 days). Given that volumetric alterations within the first 24–48 hours are expected to be modest relative to measurement variability, analyses focused on the contrast between the combined Day 0/Day 1 scans and the Day 5 scan. After quality control, 20 patients with complete longitudinal data were retained (Supplementary Table 2).

### MRI acquisition

All stroke participants were scanned using a 3T MRI systems. Acquisition parameters for both stroke cohorts are provided in the Supplementary Materials.

### Lesion segmentation

For the primary longitudinal cohort, T1-weighted MPRAGE images were used for grey matter volume (GMV) estimation, whereas lesion identification was based on multimodal structural imaging (T1, T2, and FLAIR). Lesions were manually segmented in native space using baseline structural images acquired within 2 weeks post-stroke. All segmentations were reviewed by an experienced neurologist to verify lesion boundaries and lacunar involvement. Lesion masks were subsequently normalized to MNI space.

For the early-phase cohort, lesions were manually delineated on Day 0 diffusion-weighted images. Masks were drawn by one investigator and independently verified by a second stroke imaging specialist. Lesion masks were then normalized to MNI space and dilated by one voxel (3×3×3 kernel) using FSL to mitigate potential registration mismatch.^33^ Processing scripts for this cohort are publicly available (https://github.com/sheyma/stroke_preprop).

### Longitudinal voxel-based morphometry

Longitudinal structural changes were assessed using the longitudinal VBM toolbox implemented in SPM12 (http://www.fil.ion.ucl.ac.uk/spm). For each participant, an unbiased within-subject average image was generated using the Serial Longitudinal Registration toolbox.^34^ The average images were bias-corrected and segmented into grey matter, white matter, and cerebrospinal fluid probability maps, and subsequently normalized to MNI space.

Stroke lesions were masked during segmentation and spatial normalization to prevent tissue misclassification.^35^ GM maps were modulated using Jacobian determinants derived from both native-to-average and average-to-MNI transformations to preserve volumetric information. The resulting GMV maps were smoothed using an 8-mm full-width-at-half-maximum Gaussian kernel and restricted to grey matter using a high-resolution GM mask derived from the Human Connectome Project. All preprocessing steps underwent visual quality control.

### Spatial mapping of longitudinal GMV change

After generating whole-brain voxel-wise GMV maps for all available time points, longitudinal GMV change was quantified at the individual level relative to a cohort-specific baseline. For the primary longitudinal cohort, the 2-week scan served as baseline and proportional GMV change was calculated for the 3- and 12-month scans. For the early-phase cohort, the combined Day 0/Day 1 scan served as baseline and proportional GMV change was calculated for Day 5. Proportional GMV change was defined as follows:

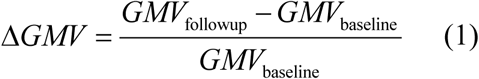

This normalization reduces potential confounding arising from regional differences in baseline GMV magnitude and is widely used in longitudinal morphometric studies of neurodegeneration and brain injury.^36,37^

Voxel-wise GMV change maps were parcellated into 384 cortical and subcortical regions using the Atlas of Intrinsic Connectivity of Homotopic Areas (AICHA),^38^ enabling integration with connectome-based modelling. Regional GMV change was computed as the mean proportional change across all constituent voxels, yielding a 384 × 1 regional GMV change vector for each subject and each time interval. All analyses were also replicated using the Brainnetome Atlas (BNA),^39^ and the corresponding validation results are reported in the Supplementary Materials.

### Normative connectome construction

Normative connectomes were derived from 930 healthy adult participants from the HCP young adult cohort, providing a reference independent of our stroke corhorts.^40,41^ The architecture of resulting networks are visualised in Supplementary Fig. 1.

Normative structural and functional connectomes were estimated from diffusion-weighted imaging and resting-state functional MRI data. All imaging data were processed using the standardized HCP minimal preprocessing pipelines,^40^ with additional denoising and quality-control procedures described in the Supplementary Materials. The resulting connectomes served as cohort-independent normative references for all connectome-based modelling analyses.

Whole-brain probabilistic tractography with anatomical constraints was performed to construct the structural connectome. The AICHA and BNA atlases were aligned to individual native diffusion space, and streamlines were assigned to region pairs based on the proximity of their endpoints to the parcellated regions. Four edge-weighting schemes were examined: binary adjacency, fractional anisotropy (FA), inverse fibre length (invFL), and fraction of streamlines (FSe). FSe was defined as the proportion of streamlines connecting two regions relative to the total number of streamlines extrinsic to those regions.^42,43^ Edges that were not reliably present across participants were excluded to reduce the influence of potentially spurious tractography-derived connections. The remaining individual weighted connectivity matrices were then averaged across participants to generate the group-level normative structural connectome.

The same AICHA and BNA atlases were used to define regional nodes, ensuring direct comparability with the structural connectome. Representative time series were extracted by averaging voxel-wise signals within each region. Functional connectivity was quantified using Pearson correlation between regional time series and subsequently Fisher z-transformed. Individual functional connectivity matrices were averaged across participants to generate the group-level normative functional connectome.

### Coordinated deformation modelling

We employed coordinated deformation modelling (CDM) to test whether longitudinal GMV change after stroke is constrained by brain network architecture. CDM was originally developed to examine whether patterns of structural deformation align with underlying connectivity structure.^44^ The model assumes that regional structural change reflects coordinated influences from network-connected regions, rather than independent or purely local processes.

Within this framework, the GMV change of each region was modelled as a weighted aggregation of GMV changes in its network neighbours, where weights correspond to inter-regional connectivity strength (Fig. 1C). CDM has previously been applied to neurodegenerative and psychiatric conditions to evaluate whether spatial patterns of group-level atrophy conform to network constraints,^24,28,45^ but it has not been systematically examined in focal brain lesions with longitudinal individual-level data.

**Figure 1.**
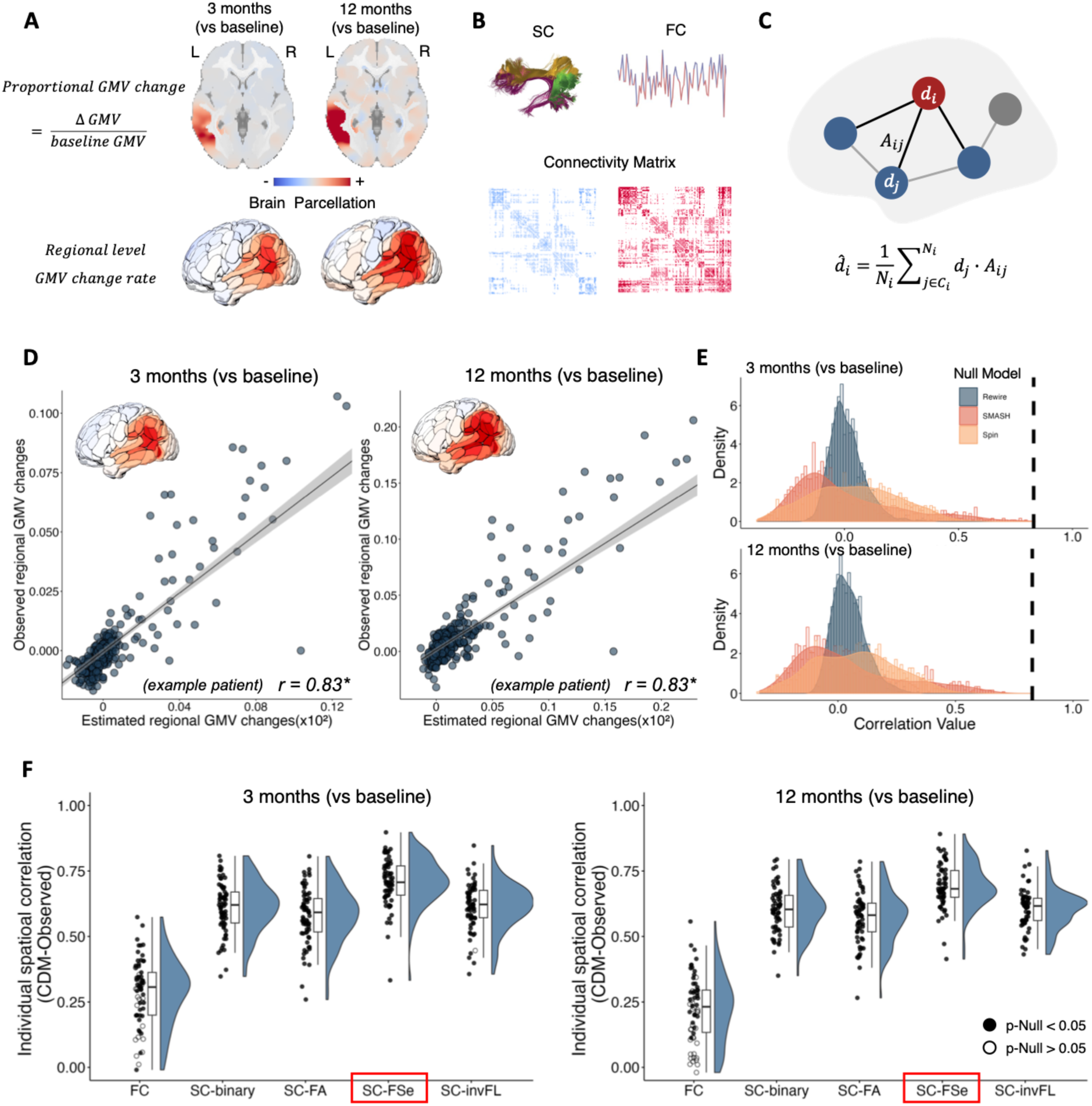
Structural connectivity constrains post-stroke longitudinal GMV atrophy. (A) Representative patient illustrating individualized proportional GMV change maps (ΔGMV / baseline GMV) at 3 and 12 months post-stroke (baseline ≈ 2 weeks). (B) Schematic of structural and functional connectome construction. Structural connectivity (SC) reflects white matter fibre pathways reconstructed from diffusion tractography using multiple edge-weighting schemes, including binary adjacency, fraction of streamlines (FSe), fractional anisotropy (FA) and inverse fibre length (invFL). Functional connectivity (FC) reflects temporal correlations of resting-state BOLD signals between regions. (C) Coordinated deformation model (CDM). Regional GMV change (*d_i_*) is modelled as the weighted aggregation of GMV changes in network-connected regions, scaled by the connectivity strength *A_ij_*. (D) Representative patient showing correspondence between CDM-estimated and empirically observed regional GMV changes at 3 and 12 months. Each point represents a brain region; lines indicate linear fits. (E) Null distributions of CDM-observation correlations generated using spatial autocorrelation-preserving, spin-based, and topology-preserving rewiring procedures. The black dashed line indicates the observed CDM correlation for the representative patient. (F) Spatial correlation between CDM-estimated and observed GMV changes across all patients. \**P* < 0.05.

Formally, for each region *i*, the estimated GMV change 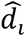 was defined as:

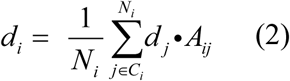

where *d_j_* denotes the observed GMV change in region *j*, *A_ij_* represents the connectivity strength between regions *i* and *j*, *C_i_* denotes the set of regions connected to *i*, and *N_i_* is the number of such connections.

CDMs were constructed using both structural and functional connectomes under multiple weighting schemes. For each subject, CDM-derived GMV change maps were compared with empirically observed maps across all available longitudinal intervals. These included the 2-week-to-3-month and 2-week-to-12-month intervals in the primary longitudinal cohort and the first week after stroke in the independent early-phase cohort. Model performance was quantified using Pearson correlation, with statistical significance assessed using spatial and network-based null models.

To determine which connectome definition best captured coordinated post-stroke GMV change across regions, we compared CDM performance across the five connectome definitions at the group level using a one-way repeated-measures ANOVA. Significant main effects were followed by Bonferroni-corrected post hoc paired t-tests.

### Network diffusion modelling

To explicitly model the dynamic propagation of post-stroke GMV change, we implemented network diffusion modelling (NDM), a generative graph-based framework for network-constrained pathological propagation (Fig. 2A-B).^46,47^

**Figure 2.**
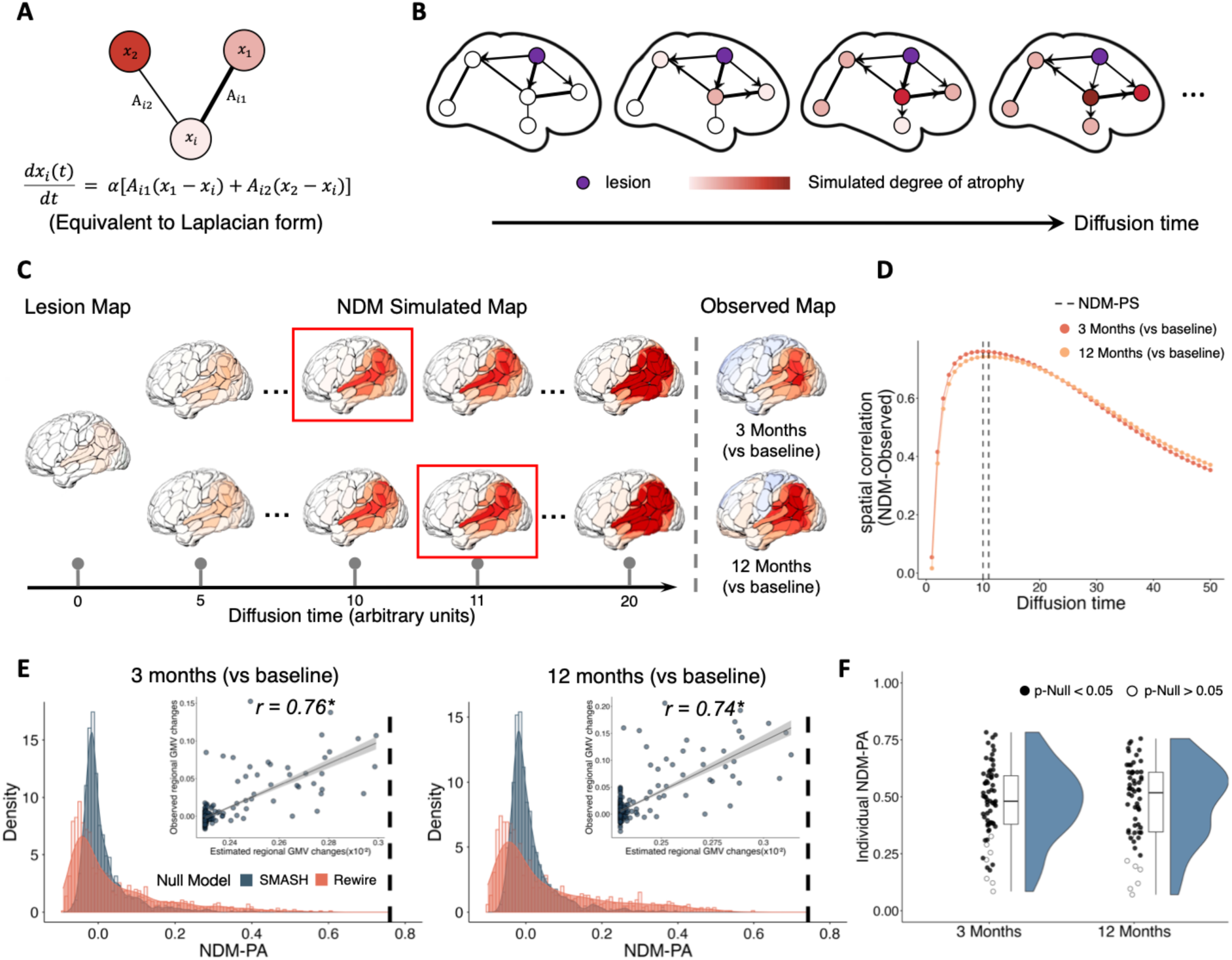
Network diffusion modelling captures lesion-driven propagation of GMV atrophy. (A) Network diffusion model describing regional change as a function of weighted differences between each region and its connected neighbours, with coupling strengths defined by the connectivity weight. (B) Conceptual illustration of lesion-initiated propagation of pathological load through the structural connectome over diffusion time. Diffusion time is expressed in arbitrary units and does not correspond to chronological time. (C) Representative patient illustrating lesion location, network diffusion modelling (NDM)-simulated GMV atrophy maps across diffusion time, and empirically observed GMV atrophy maps at 3 and 12 months post-stroke. For each observation, the NDM-simulated map achieving the highest spatial correlation with the empirical GMV atrophy pattern is highlighted by the red rectangle. (D) Spatial correlation between NDM-predicted and observed GMV atrophy maps as a function of diffusion time for the representative patient. The maximum spatial correlation achieved across diffusion time was defined as the NDM prediction accuracy (NDM-PA), and the corresponding diffusion time was defined as the NDM propagation stage (NDM-PS) for that sampling time point. (E) Null distributions of NDM-PA generated using spatial autocorrelation-preserving, spin-based, and topology-preserving rewiring procedures. The dashed line indicates the observed NDM-PA for the representative patient. Insets show correspondence between predicted and observed regional GMV changes at the NDM-PS. (F) Distributions of individual NDM-PA values across all patients at 3 and 12 months post-stroke. Filled symbols indicate patients exceeding null thresholds. \**P* < 0.05.

NDM formalizes pathological spread as a first-order diffusion process on the graph Laplacian of a connectivity matrix. The evolution of the regional pathological state vector *X*(*t*) is governed by:

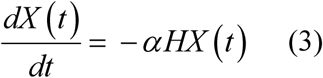

where *X*(*t*) represents the simulated pathological load across regions at diffusion time *t*, *H* represents the graph Laplacian of the normative connectivity matrix, and *⍺* is a global diffusion rate parameter fixed to 1 for all subjects.

The closed-form solution is:

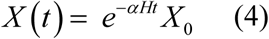

where *X*_0_ specifies the initial pathological state. In this study, the binary lesion map served as *X*_0_, reflecting the hypothesis that secondary degeneration originates from the focal injury site and propagates along connectome-defined pathways. Diffusion time *t* is expressed in arbitrary units and does not correspond directly to chronological time.

For each subject, NDM simulations were performed across a continuous diffusion-time interval (*t* ∈ [0,50]), sufficient to capture the full range of propagation dynamics (Fig. 2C). At each diffusion time point, the simulated GMV change pattern was compared with the empirically observed GMV change map using Pearson correlation (Fig. 2D). The maximum correlation value across diffusion time was defined as the NDM prediction accuracy (NDM-PA), and the statistical significance of the NDM-PA value was evaluated using spatial null models (Fig. 2E).

The diffusion time at which this maximum correlation was achieved corresponds to the model-inferred propagation stage that best matches the observed GMV change pattern. This time point was defined as the NDM propagation stage (NDM-PS) for that sampling time point post-stroke.

### Null models

Multiple complementary null models were implemented to determine whether CDM or NDM performance exceeded that expected under spatial and topological constraints.

Spatial autocorrelation-preserving surrogate maps were generated using the BrainSMASH framework,^48^ which retains the empirical spatial autocorrelation structure while permuting regional values. In addition, spin-based nulls were applied to control for cortical spatial embedding.^48,49^ Intrinsic network topology was addressed through degree-preserving random rewiring implemented in the Brain Connectivity Toolbox,^50^ generating topology-matched surrogate connectomes.

CDM analyses incorporated spatial autocorrelation-preserving, spin-based, and topology-preserving nulls, whereas NDM analyses utilized spatial autocorrelation and topology-preserving nulls. Detailed implementation procedures are provided in the Supplementary Materials.

### Within-subject comparison of chronological time and NDM propagation stage

The relationship between chronological progression after stroke and model-inferred propagation stage was examined by comparing NDM-PS values derived from GMV change patterns at 3 and 12 months within subjects with complete longitudinal imaging. Group-level differences in NDM-PS between the two time points were assessed using paired t-tests. Within-subject changes were quantified as ΔNDM-PS = NDM-PS_12 months_ − NDM-PS_3 months_.

This longitudinal quantification enabled the stratification of subjects according to whether chronological ordering from 3 to 12 months was concordant with diffusion stage: subjects exhibiting an increase in propagation stage over chronological time were defined as the concordant group (NDM-PS_12 months_ > NDM-PS_3 months_), whereas the others were classified as the discordant group (NDM-PS_12 months_ ≤ NDM-PS_3 months_).

Spatial similarity between 3 and 12-month GMV change maps was quantified using Pearson correlation. In addition, an interval-specific GMV change map (3-to-12 months) was constructed for each subject to characterize incremental structural change during the chronic stage. NDM simulations were further applied to these incremental GMV change maps, and NDM-PA was compared between concordant and discordant groups.

### Multi-level correlates of NDM parameters

Following the observation of heterogeneous longitudinal propagation-stage shifts at 12 months, analyses of inter-individual correlates of NDM-derived parameters were conducted at the 3-month post-stroke time point, at which propagation-stage estimates remained more comparable across subjects.

We first examined whether demographic variables, including age, sex, handedness, and years of education, were associated with NDM-PA and NDM-PS.

We then examined lesion-related effects across multiple levels, including lesion topography, global lesion characteristics, and the network-topological and molecular properties of lesioned regions. At the regional level, individual NDM parameters were mapped onto lesioned regions defined by the AICHA and BNA atlases and averaged across subjects with lesions involving each region, generating region-wise NDM-PA and NDM-PS maps. Regions with higher mean NDM-PA or NDM-PS indicated locations where lesion involvement was associated with better NDM performance or later inferred propagation stage, respectively. System-level effects were assessed by assigning region-wise NDM parameter maps to canonical functional systems and testing for between-system differences in parameter values. Between-system differences were evaluated using a Kruskal–Wallis test,^51^ followed by Bonferroni-corrected two-sided Mann–Whitney U tests for pairwise post hoc comparisons.^52,53^

At the subject level, lesion hemisphere and lesion volume were tested for associations with NDM-PA and NDM-PS across subjects. To evaluate the role of lesion-site network topology, nodal properties of AICHA- and BNA-defined regions—including degree centrality, betweenness centrality, local clustering coefficient, and nodal efficiency—were derived from the normative structural connectome above. Subject-specific topological features were obtained by averaging these nodal metrics across all lesioned regions and were then tested for associations with NDM-PA and NDM-PS across subjects. In addition, to assess the contribution of the molecular environment of lesioned regions, neurotransmitter receptor and transporter density maps were obtained using the neuromaps toolbox.^54,55^ For each subject, receptor/transporter densities were averaged across all lesioned regions to generate subject-specific molecular profiles, and their associations with NDM-PA and NDM-PS were evaluated across subjects. Statistical significance for all subject-level associations was determined using permutation tests by shuffling the NDM metrics across subjects to generate null distributions.^56^ The resulting p-values were then corrected for multiple comparisons within each analysis category using the false discovery rate (FDR) method, with significance set at *P_FDR_* < 0.05.^57^

### NDM-based prediction for individualized GMV atrophy patterns

Building on the NDM framework described above, we next tested whether lesion topology contains sufficient information to predict downstream structural degeneration at the individual level. A schematic overview of this predictive workflow is presented in Fig. 5A.

Ridge regression was used to estimate subject-specific NDM propagation stage (NDM-PS) from lesion-derived spatial features (Fig. 5B).^58^ L2 regularization was applied to stabilize parameter estimation in the high-dimensional feature space. Model training and evaluation were conducted using a nested leave-one-out cross-validation framework, in which one subject was held out for testing in each outer fold and the remaining subjects were used for training. An inner cross-validation loop was used to optimize the regularization parameter.^59^

Predicted NDM-PS values were incorporated into NDM to generate individualized simulated GMV atrophy maps. Prediction performance was quantified as the spatial correlation between simulated and empirically observed GMV atrophy patterns. Statistical significance was evaluated using permutation testing within the cross-validation framework.^60^

This procedure enabled evaluation of whether lesion-informed NDM-PS estimation supports individualized prediction of longitudinal structural degeneration patterns without reliance on longitudinal imaging data.

## Results

We analysed longitudinal MRI data from a primary cohort (n=72, age: 53.9±10.5 years) and an independent early-phase cohort (n=20, age: 63.3±12.03 years), spanning hyperacute to chronic stages. Detailed demographic and clinical information for all participants is available in Supplementary Tables 1 and 2.

### Structural connectivity constrains post-stroke GMV atrophy

Across patients, longitudinal GMV changes at 3 and 12 months post-stroke exhibited spatially coordinated patterns extending beyond focal lesion regions (Fig. 1A). CDM-estimated maps showed strong spatial correspondence with observed GMV changes at both time points (Fig. 1D). CDM performance varied across connectome definitions, with structural connectivity-based models consistently outperforming functional connectivity-based models at both 3 and 12 months (Fig. 1F; structural connectivity, *r* range: 0.26–0.90; functional connectivity, *r* range: −0.02–0.57). Repeated-measures ANOVA confirmed differences across connectome definitions at both time points (both *P* < 0.001). Post hoc paired comparisons showed that functional connectivity yielded lower CDM–observation correspondence than all four structural connectome definitions, whereas FSe-weighted connectomes yielded higher correspondence than all other definitions (FSe-weighted: *r* range: 0.33–0.90; mean *r* [SD] = 0.71 [0.09] at 3 months and 0.69 [0.09] at 12 months). Most patients showed significant CDM–observation correspondence. The same overall pattern was replicated under the BNA parcellation (Supplementary Fig. S2A-C).

Applying the same analysis to the early-phase cohort spanning the hyperacute and early acute stages within the first week after stroke revealed a similar overall pattern. Although early GMV changes were more spatially diffuse and less lesion-centred (Supplementary Fig. S3A), structural connectivity-based CDMs again outperformed functional connectivity–based models, with FSe-weighted connectome again yielding the strongest correspondence (mean *r* [SD] = 0.50 [0.13]; Supplementary Fig. S3B). Differences across connectome definitions were again present (*P* < 0.001), with functional connectivity yielding the lowest correspondence overall and FSe-weighted connectomes yielding the highest. Effect sizes were reduced relative to later stages, but structural connectivity constraints were already detectable within the first week after stroke. Together, these findings indicate that post-stroke atrophy follows structural connectome constraints from the hyperacute to chronic stages.

### Network diffusion modelling captures lesion-driven propagation

Building on the identification of structural connectivity constraints in coordinated GMV atrophy, we applied NDM using the FSe-weighted structural connectome to simulate lesion-initiated propagation of GMV atrophy at the individual level (Fig. 2A-B). Using each patient’s lesion profile as the initial condition, NDM generated time-resolved predictions of distributed GMV change patterns extending beyond focal lesion sites. An example patient illustrating this workflow is shown in Fig. 2C-E.

Across patients, NDM-PA values were consistently positive at both 3 and 12 months post-stroke (mean *r* [SD] = 0.47 [0.16] at 3 months; 0.48 [0.17] at 12 months), with 89.9% and 87.1% of patients, respectively, exhibiting significant NDM-PA values (Fig. 2F). These results were replicated using the BNA parcellation (Supplementary Fig. S2D). Together, these findings indicate that NDM captures lesion-driven, patient-specific patterns of longitudinal GMV atrophy.

This pattern was not yet fully expressed in the early-phase dataset within the first week after stroke. At this stage, NDM-PA values were markedly attenuated, and most patients (95%) failed to reach significance (Supplementary Fig. S3C). This contrast suggests that lesion-initiated, network-consistent atrophy trajectories become progressively organized across subacute and chronic stages rather than being fully established immediately after stroke.

### Chronological time after stroke does not consistently track NDM propagation stage

If post-stroke structural degeneration were aligned with network diffusion dynamics, later chronological time points (12 months) would correspond to later model-inferred propagation stages (NDM-PS) within individuals. However, NDM-PS values did not differ between 3 and 12 months at the group level (paired t-test: *t*(58) = 0.47, *P* > 0.05, two-tailed; Fig. 3A). Within-subject shifts in propagation stage (ΔNDM-PS = NDM-PS_12 months_ − NDM-PS_3 months_) were broadly distributed around zero (Fig. 3A-B), indicating substantial inter-individual heterogeneity.

**Figure 3.**
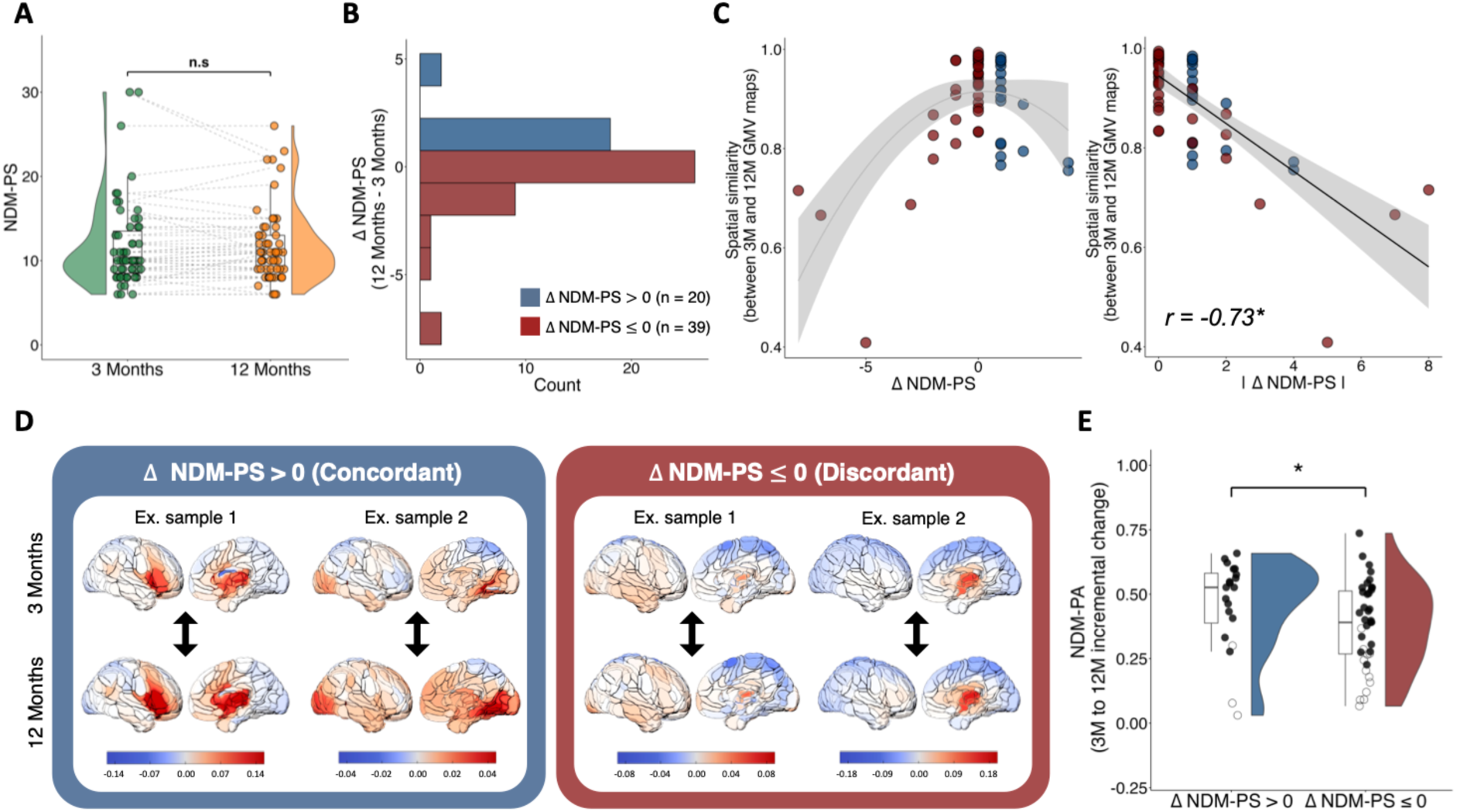
Chronological time after stroke does not consistently track NDM propagation stage. (A) Comparison of NDM propagation stage (NDM-PS) estimated from GMV atrophy patterns at 3 and 12 months post-stroke. (B) Distribution of within-subject differences in propagation stage (ΔNDM-PS = NDM-PS_12months_ -NDM-PS_3months_). (C) Association between propagation-stage shifts and empirical spatial reorganization of atrophy. Left: spatial similarity between observed 3- and 12-month GMV maps as a function of ΔNDM-PS. Right: larger absolute stage shifts (|ΔNDM-PS|) are associated with lower spatial similarity. (D) Representative examples of patients stratified by chronological-stage ordering concordance (concordant or discordant). (E) NDM prediction accuracy (NDM-PA) for incremental 3-to-12-month GMV change maps for the concordant and discordant patient groups (one-tailed independent sample *t* test). n.s: non-significant, \**P* < 0.05.

We then examined the association between ΔNDM-PS and empirical pattern similarity to determine whether this time–stage dissociation reflects metric instability or meaningful variation in longitudinal atrophy organization. Larger absolute shifts in propagation stage (|ΔNDM-PS|) were associated with lower spatial similarity between observed 3- and 12-month GMV change maps (Pearson’s *r* = −0.73, *P* < 0.001; Fig. 3C), indicating that variation in NDM-PS corresponds to differences in spatial reorganization of atrophy.

Patients were then stratified according to whether chronological ordering (3 months < 12 months) was concordant with propagation-stage ordering. Twenty subjects exhibited concordant progression (ΔNDM-PS > 0), whereas thirty-nine showed discordant ordering (ΔNDM-PS ≤ 0). As shown in Fig. 3D, the concordant group displayed more spatially extensive or intensified GMV atrophy at 12 months relative to 3 months, whereas the discordant group showed largely overlapping patterns across time. Consistent with this contrast, NDM simulations applied to the incremental 3-to-12-month GMV change map yielded higher NDM-PA in the concordant group than in the discordant group (two-sample t-test: *t*(57) = 1.79, *P* < 0.05, one-tailed; Fig. 3E). These findings suggest that advancement in NDM propagation stage is associated with continued network-consistent atrophy evolution, whereas a substantial subset of patients diverges from diffusion-consistent propagation during the chronic stage. Validation analyses using the BNA yielded highly comparable overall trends, with the core patterns of atrophy evolution remaining stable across both templates (Supplementary Fig. S4).

### Lesion-related factors are associated with NDM parameters at 3 months post-stroke

Given the heterogeneous longitudinal shifts in NDM propagation stage observed at 12 months, we focused on the 3-month post-stroke time point, at which NDM-PS remained more comparable across subjects, and examined multi-level lesion-related factors associated with inter-individual variability in NDM-derived parameters.

Neither NDM-PA nor NDM-PS showed significant associations with demographic variables, including age, sex, handedness, and years of education. In contrast, lesion-related factors exhibited clear multi-level associations with NDM parameters.

At the level of lesion topography, we mapped NDM parameters onto lesioned regions and averaged them across subjects with lesions involving each region, revealing marked spatial specificity (Fig. 4A-B). Regions whose lesion involvement was associated with higher mean NDM-PA were concentrated in the left precuneus, posterior cingulate cortex, and superior parietal lobule, indicating that lesions affecting these regions were associated with more predictable atrophy patterns. Regions associated with higher mean NDM-PS included the left parieto-occipital gyrus, right thalamus, and left anterior cingulate cortex, indicating later inferred propagation stages when these regions were involved. Consistent with this spatial heterogeneity, NDM-PS differed significantly across canonical functional systems (Kruskal–Wallis *H* = 85.81, *P* < 0.001), with subcortical and somatomotor systems showing higher propagation stages than other systems, whereas NDM-PA showed no significant between-system differences (Fig. 4C).

**Figure 4.**
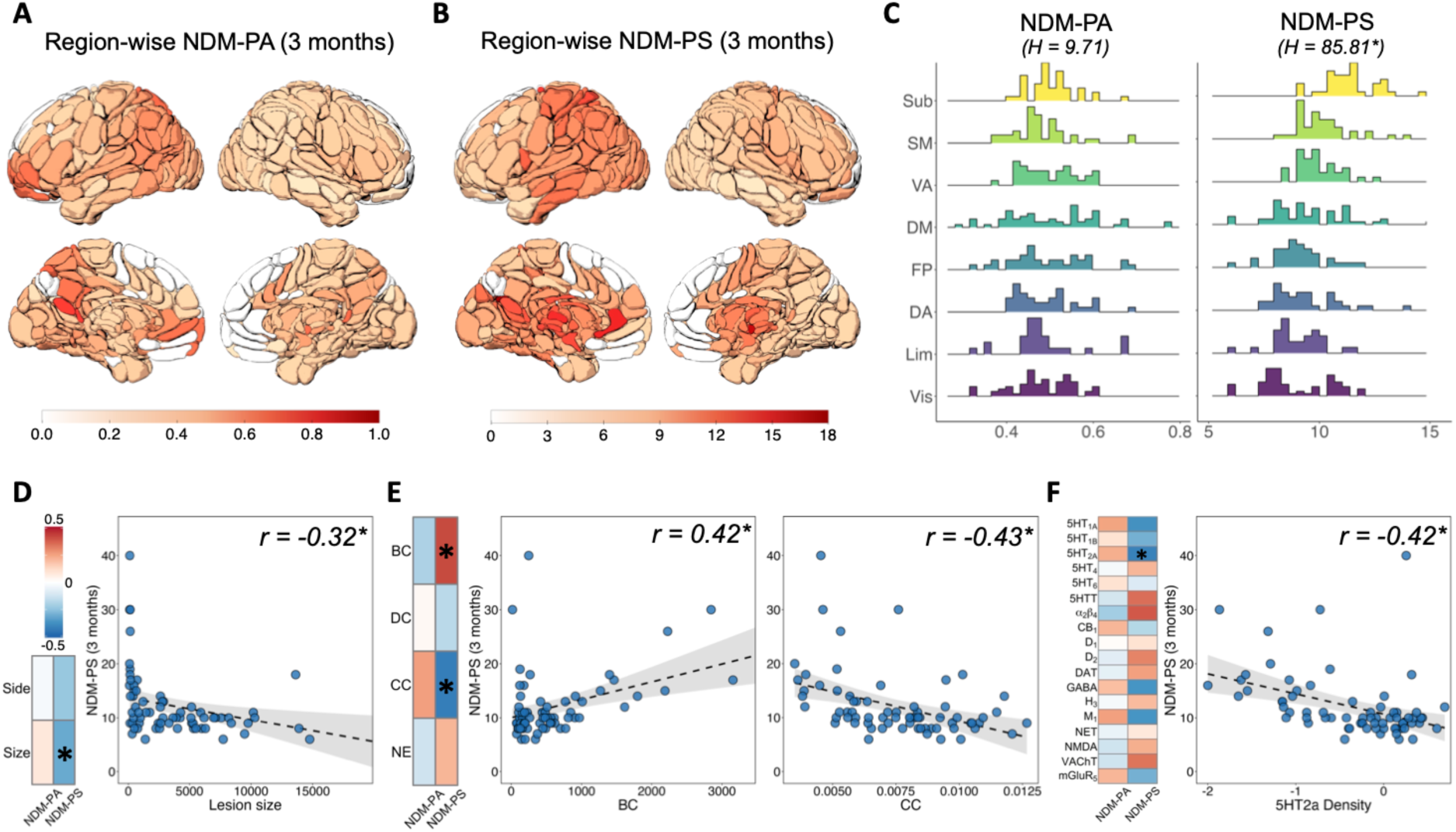
Multi-level lesion-related associations of NDM parameters at 3 months post-stroke. (A) Region-wise map of NDM-PA at 3 months post-stroke. Values indicate the mean NDM-PA across subjects with lesions involving the corresponding region. (B) Region-wise map of NDM-PS at 3 months post-stroke. Values indicate the mean NDM-PS across subjects with lesions involving the corresponding region. (C) Distribution of region-wise NDM-PA and NDM-PS across the seven Yeo functional networks and a subcortical system. NDM-PS differed significantly across systems (Kruskal–Wallis test: *H* = 85.81, *P* < 0.001), whereas NDM-PA did not (*H* = 9.71, *P* > 0.05). Vis, visual; SM, somatomotor; DA, dorsal attention; VA, ventral attention; Lim, limbic; FP, frontoparietal; DM, default mode; Sub, subcortical. (D-F) Subject-level associations of NDM-PA and NDM-PS with lesion-related factors. Heatmaps show Pearson correlation coefficients, and scatterplots show associations surviving permutation-based false discovery rate correction. Dashed lines and shaded bands indicate linear fits and 95% confidence intervals. Asterisks indicate associations surviving FDR correction (*P*_FDR_ < 0.05). (D) Subject-level associations of NDM-PA and NDM-PS with global lesion characteristics, including lesion side and lesion size. (E) Subject-level associations of NDM-PA and NDM-PS with network-topological properties of lesioned regions, including betweenness centrality (BC), degree centrality (DC), clustering coefficient (CC), and nodal efficiency (NE). (F) Subject-level associations of NDM-PA and NDM-PS with neurotransmitter receptor/transporter profiles of lesioned regions derived from PET maps.

At the individual level, global lesion characteristics showed selective associations with NDM parameters. NDM-PA was not significantly related to lesion hemisphere or lesion size. By contrast, NDM-PS decreased with increasing lesion size (*r* = −0.32, *P_FDR_* < 0.05), indicating that larger lesions were associated with less advanced propagation stages at 3 months (Fig. 4D).

The network-topological and molecular properties of lesioned regions were also differentially related to NDM parameters. NDM-PA showed no significant associations with lesion-site topological or molecular features. In contrast, NDM-PS exhibited a broader association pattern: longer propagation stages were associated with higher betweenness centrality of lesioned regions (*r* = 0.42, *P_FDR_* < 0.05), whereas shorter propagation stages were associated with higher local clustering coefficient (*r* = −0.43, *P_FDR_* < 0.05) (Fig. 4E). At the molecular level, NDM-PS was additionally associated with neurotransmitter receptor/transporter profiles of lesioned regions, with the strongest effect observed for the 5-HT2a receptor (*r* = −0.42, *P_FDR_* < 0.05) (Fig. 4F). Together, these findings indicate that variation in NDM propagation stage is shaped by lesion topography, lesion size, and the network-topological and molecular context of the damaged region, whereas NDM prediction accuracy is comparatively less sensitive to these subject-level determinants.

Results were broadly consistent under the BNA parcellation, supporting the robustness of these associations across atlas definitions (Supplementary Fig. S5).

### Lesion information supports individualized prediction of GMV atrophy

We next evaluated whether lesion information alone is sufficient to predict individualized GMV atrophy patterns within the NDM framework. For each patient, lesion maps were used as input to NDM to generate time-resolved simulated GMV atrophy patterns. To enable prediction without longitudinal imaging, a regression model was trained to estimate subject-specific NDM-PS from lesion-derived spatial features using leave-one-out cross-validation (Fig. 5A–B).

**Figure 5.**
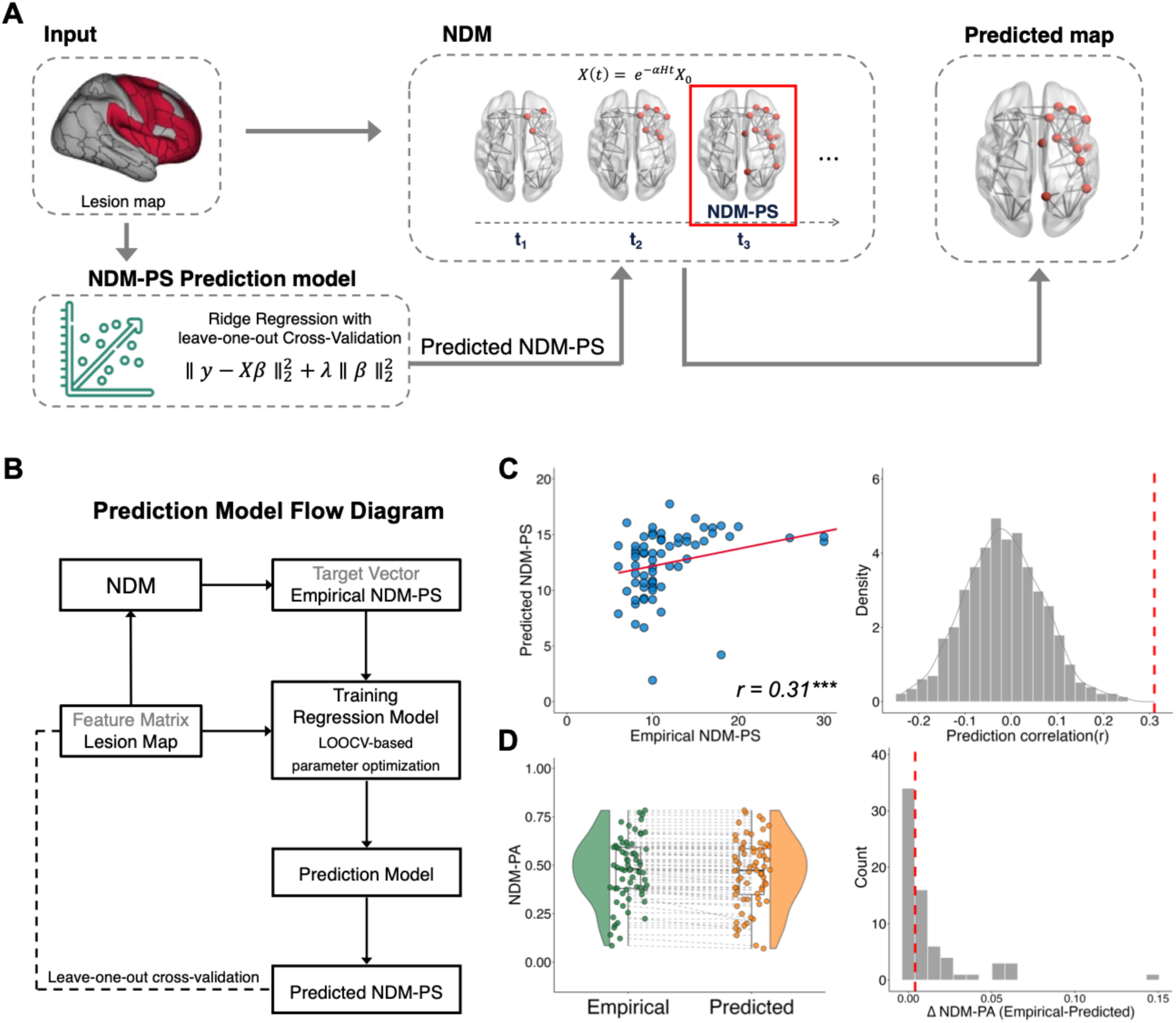
Lesion-informed NDM enables individualized prediction of GMV atrophy. (A) Framework for individualized prediction of GMV atrophy patterns. Lesion maps serve as input to NDM to generate time-resolved simulated GMV change maps. A ridge regression model with leave-one-out cross-validation is trained to predict subject-specific NDM-PS from lesion-derived features. The predicted NDM-PS is then used to select the corresponding NDM-simulated GMV map for each individual. (B) Flow diagram of the prediction procedure. Lesion-derived regional features are used to train a regression model to estimate empirical NDM-PS values. The trained model generates out-of-sample predicted NDM-PS for each subject, which are subsequently input to NDM to produce individualized GMV atrophy predictions. (C) Prediction accuracy of NDM-PS. Left: correlation between predicted and empirically derived NDM-PS values across individuals. Right: null distribution of prediction correlations obtained from permutation testing; the dashed line indicates the observed prediction performance. (D) Spatial correspondence between observed GMV atrophy patterns and NDM predictions generated using empirically derived NDM-PS versus regression-predicted NDM-PS. Group-level distributions of NDM-PA demonstrate comparable spatial performance between the two approaches. The right panel shows the distribution of differences in NDM-PA between empirical-stage and predicted-stage models. \*\*\**P* < 0.001.

Across participants, regression-predicted NDM-PS values were significantly correlated with empirically derived NDM-PS values obtained by aligning NDM simulations to observed longitudinal GMV atrophy patterns (*r* = 0.31, *P* < 0.001; Fig. 5C), exceeding permutation-based null expectations.

Using the predicted NDM-PS values, individualized NDM-based GMV atrophy maps were generated. At the group level, predicted maps demonstrated spatial correspondence with observed GMV changes comparable to those generated using empirically derived NDM-PS values (Fig. 5D). Similar predictive performance was observed under the BNA parcellation (Supplementary Fig. S6).

Together, these findings suggest that lesion-informed estimation of NDM propagation stage enables individualized prediction of GMV atrophy patterns without requiring longitudinal imaging data.

## Discussion

By leveraging two complementary longitudinal cohorts spanning the hyperacute to late chronic stages after stroke, the present study shows that post-stroke atrophy follows individual-level trajectories that are strongly shaped by structural brain networks. Across analyses, a coherent picture emerged: remote atrophy was not distributed arbitrarily across the brain, but was organized by the structural connectome; later atrophy patterns were not only network dependent in a generic sense, but were linked to each patient’s acute lesion as an initial condition; and the spatiotemporal expression of this process varied systematically with the topological and molecular context of the lesion. Together, these findings suggest that focal stroke initiates a distributed pattern of secondary alternation whose spatial evolution is shaped by large-scale structural brain architecture.

Our results identified the structural connectome as the dominant scaffold for longitudinal post-stroke degeneration. CDM analyses showed that regions tended to exhibit coordinated GMV loss with their structurally connected neighbours, and that this dependence was already detectable in the hyper-acute stage before becoming more pronounced over time. By contrast, functional connectivity provided a much weaker account of coordinated change. This contrast suggests that the large-scale organization of remote degeneration is anchored less in transient functional coupling^61–63^ than in the relatively stable anatomical pathways^64^ through which lesion effects are embedded and distributed.^65,66^ In this respect, the coordinated decline observed via CDM might reflects biological processes mediated by the underlying white matter, whereby Wallerian degeneration and inflammatory signalling propagate through anatomical links to synchronise structural loss across connected regions.^67–71^ Our findings support a shared principle across neurological and psychiatric conditions, where the structural connectome acts as a universal template constraining how diverse perturbations are translated into distributed structural abnormalities.^72–75^

The superior performance of the FSe-weighted connectome underscores this point further. Binary connection presence alone, as well as weights derived from path length or microstructural metrics, proved less predictive than a representation that also captured the relative strength or capacity of anatomical projections^42^. This suggests that what matters is not merely the existence or distance of a pathway, but the degree to which a region is embedded within the white matter architecture that scales the transmission of lesion effects.^76^ More broadly, this finding raises the possibility that network models of post-stroke degeneration should be sensitive not simply to topology, but to graded differences in the capacity of structural pathways to support coordinated downstream change.

If CDM identifies structural constraint, NDM extends that observation into a more specific hypothesis about process. When initialized with each patient’s acute lesion map, NDM generated simulated atrophy patterns that significantly corresponded to observed GMV changes at both 3 and 12 months post-stroke. Under this framework, post-stroke structural reorganization can be well captured by a network-mediated process in which lesion location serves as a patient-specific initial condition, and subsequent changes emerge over time as a function of each region’s structural coupling to affected regions.^46,47^ The simulated dynamics of the NDM may find a plausible biological basis in transneuronal degeneration, a potential driver of secondary decay where neurons undergo deterioration following the loss of afferent input or trophic support.^77–81^ Such an interpretation is supported by prior observations that regions structurally connected to the lesion are more likely to exhibit subsequent atrophy.^76,82,83^ Under this view, the diffusion-like behaviour captured by NDM is best interpreted not as literal spread of pathogenic agents, but as a useful description of how connection-dependent vulnerability becomes expressed across the connectome after focal injury. Notably, the robust performance of the NDM across diverse lesion configurations—despite substantial heterogeneity in size and location—strengthens the argument that the structural connectome imposes a canonical organizing principle on post-stroke degeneration. In this light, clinical heterogeneity serves not as confounding noise, but as a critical perturbation that confirms the generalizability of our network-based framework.

Furthermore, the temporal profile of the NDM results highlights that this network process is not fully expressed immediately after stroke. Model performance remained limited in the early-acute stage, even though structural constraint was already detectable with CDM. This divergence suggests that early network dependence and subsequent network-consistent propagation is related but not identical features of post-stroke degeneration. One implication is that focal injury may first establish the conditions for distributed vulnerability, whereas the transnueronal cascades modelled by the NDM require weeks or months to manifest, ultimately driving a diffusion-like pattern of macroscopic volume loss.^84–88^ The present NDM should therefore be viewed as distilling one dominant organizing principle of later degeneration, rather than an exhaustive account of the diverse pathophysiology processes present across the entire post-stroke time course.^89–91^

This perspective also helps clarify the meaning of NDM-derived propagation stage in the context of longitudinal atrophy evolution. The present results suggest that propagation stage should not be treated as a simple proxy for chronological time after stroke. Although NDM captured atrophy patterns at both 3 and 12 months, inferred propagation stage did not increase monotonically across individuals. Instead, it reflects the topological depth of the diffusive progression, marking the specific point in the simulated diffusion process where network-mediated patterns best align with empirical observations. At the group level, the limited advancement from 3 to 12 months raises the possibility that the major topographic organization of secondary degeneration is established relatively early supported by the high spatial stability of atrophy patterns observed between these sessions. This stagnation likely reflects a biological plateau in the subacute-to-chronic transition, where the neural plasticity or intrinsic resistance mechanisms emerge to counteract further pathological propagation.^87,92,93^ NDM propagation stage therefore is better characterized as a summary index of how closely observed atrophy conforms to a network-consistent diffusion-like pattern, rather than as a literal measure of elapsed biological time.

The factors associated with variation in this index further reinforce the idea that post-stroke degeneration depends on lesion embedding within a broader network context. Inter-individual variation in propagation stage was related primarily to lesion-related features rather than demographic variables. Larger lesions were associated with shorter inferred propagation stages, whereas lesions involving regions with high betweenness centrality or low local clustering showed patterns suggestive of broader network involvement. This aligns with a ‘targeted attacks’ framework, where damage to highly central elements disproportionately perturb large-scale organization.^94,95^ Furthermore, propagation stage was significantly associated with the 5-HT2a and 5-HT1a receptor density of the lesioned regions. These molecular associations are biologically plausible given convergent evidence that serotonergic signalling—particularly 5-HT1a—is implicated in neuroprotection, post-ischaemic plasticity, and functional recovery,^96,97^ whereas 5-HT2a-related variation has also been associated with ischaemic stroke susceptibility, potentially through serotonergic effects on vascular contraction and platelet aggregation.^98^ Collectively, these observations suggested that the trajectory of distributed atrophy depends on where a lesion sits within the brain’s integrated topological and molecular landscape, which collectively modulates how structural pathways are engaged following focal injury._82,83,99_

The lesion-informed prediction results also highlight the potential translational value of this framework. In contrast to prior work that either predicted future tissue loss from lesion-induced connectivity disruption or used distributed post-stroke morphometry patterns for individualized prediction,^83,100^ the proposed workflow uses acute lesion information to estimate subject-specific whole-brain GMV reorganization through the structural connectome. By incorporating the NDM, the prediction shifts from local damage alone to lesion embedding within a network model of secondary degeneration. Although still a proof of concept, this predictive mapping could facilitate clinical patient stratification by identifying those most susceptible to widespread neurodegeneration. The methodology offers a clinically relevant strategy for anticipating individual patterns of remote structural change, thereby informing tailored neuroprotective interventions or cognitive rehabilitation programs before such changes become consolidated.^101–104^

A few limitations warrant consideration. First, while the NDM effectively captures network-mediated atrophy, its simplified framework overlooks concurrent neuroplastic remodelling or compensatory hypertrophy, suggesting the integration of more biologically nuanced models in future studies. Second, as the current study utilized normative connectomes as a reference, future work with patient-specific network data may refine atrophy mapping and clarify how direct network alterations relate to structural decay; although normative functional connectome showed limited explanatory power for coordinated GMV loss here, directly measured functional connectome of patients may perform better and inform cognitive recovery after stroke.^105,106^ Finally, linking NDM-predicted atrophy to specific clinical deficits and assessing the modulatory effects of treatment remains a crucial next step to establish the translational value of this framework.

In conclusion, the present study indicates that focal stroke gives rise to individual trajectories of post-stroke atrophy that are strongly shaped by the structural connectome. This process is partially captured by a lesion-initiated, diffusion-like model, varies with the topological and molecular context of the lesion, and may provide a basis for individualized prediction of later whole-brain structural reorganization after stroke.

## Supporting information

Supplementary methods and results

## Data availability

The data supporting the findings of this study are derived from two primary sources. The main longitudinal cohort data are publicly accessible through the Central Neuroimaging Data Archives (CNDA) at https://cnda.wustl.edu/ under the reference number CCIR_00299. For the early-phase cohort, data are not publicly available due to institutional restrictions and data ownership agreements; however, data may be made available from Şeyma Bayrak at upon reasonable request, subject to a formal data use agreement and institutional approval.

