## Supplementary methods and results for "Structural brain networks shape individual-level progression of brain atrophy after stroke"

#### MRI acquisition

**Main longitudinal cohort:** MRI data for the primary longitudinal cohort were acquired using a 3T Siemens Tim-Trio scanner equipped with a 12-channel head coil at Washington University School of Medicine. The structural imaging protocol included: a sagittal T1-weighted MPRAGE sequence (TR = 1950 ms, TE = 2.26 ms, flip angle = 9°, 1.0-mm isotropic voxel size); a transverse T2-weighted turbo spin-echo sequence (TR = 2500 ms, TE = 435 ms, 1.0-mm isotropic voxel size); and a sagittal fluid-attenuated inversion recovery (FLAIR) sequence (TR = 7500 ms, TE = 326 ms, 1.5-mm isotropic voxel size).

**Independent early-phase cohort:** For the independent early-phase cohort, imaging was performed on a 3T Siemens Tim-Trio scanner following a standardized clinical protocol.[1] The acquisition included: a 3D T1-weighted MPRAGE sequence (TR = 1.9 s, TE = 2.52 s, TI = 0.9 s, 192 slices, flip angle = 9°, 1.0-mm isotropic voxel size); diffusion-weighted imaging (DWI; TR = 8.2 s, 50 volumes, 2×2×2.5 $mm^3$ voxel size); and a FLAIR sequence (TR = 8.0 s, 54 volumes, 0.5×0.5×5 $mm^3$ voxel size).

#### Normative connectome construction

##### Normative structural Connectome

Diffusion-weighted images (DWI) were processed using the HCP minimal preprocessing pipeline.[2] Voxel-wise fiber orientation distributions (FODs) were estimated via multi-shell multi-tissue constrained spherical deconvolution (CSD) with maximum harmonic order L_max = 8.[3,4] Whole-brain probabilistic tractography was performed using the second-order integration over fiber orientation distributions (iFOD2) algorithm in MRtrix3.[5] Anatomically constrained tractography (ACT) was implemented using tissue partial volume maps (PVMs) derived from T1-weighted image segmentation.[6,7] Tracking parameters were: Step size, 0.625 mm; Maximum curvature per step, 45°; FOD amplitude cutoff, 0.05; Streamline length range: 2.5–250 mm.

For each participant, 100 million streamlines were generated with seeding and termination constrained to the gray matter–white matter interface (GMWMI). The tractogram was then filtered to 10 million streamlines using the Spherical-deconvolution Informed Filtering of Tractograms (SIFT) procedure to ensure correspondence between streamline density and underlying fiber density.[8]

To construct individual structural connectomes, streamlines were assigned to the nearest atlas region within a 2-mm radius sphere centered at each streamline endpoint. AICHA and BNA parcellations were transformed to native diffusion space for streamline assignment. Four edge-weighting schemes were implemented: binary adjacency, fractional anisotropy (FA), inverse fiber length (invFL), and fraction of streamlines (FSe). In the binary scheme, edges were coded as present or absent, whereas in the other schemes they were weighted by FA, invFL, or FSe. Specifically, the FSe weight was defined as the proportion of streamlines connecting two regions relative to the total number of streamlines extrinsic to both regions (i.e., excluding intra-regional streamlines), thereby normalizing inter-regional connectivity strength for regional size and overall tractography density.[9,10]

To mitigate potential tractography-related spurious connections, only edges reliably observed across participants were retained when constructing the group-level normative structural connectome. Individual connectivity matrices were averaged to generate the final normative matrix.

**Normative functional Connectome**

Resting-state fMRI data were processed using the HCP functional minimal preprocessing pipeline.[2] Data were further denoised using ICA-FIX to remove non-neural components.[11] Subsequent preprocessing steps included: linear detrending, nuisance regression (6 head motion parameters, white matter, CSF, and global signals), band-pass filtering (0.01–0.1 Hz), spatial smoothing (6-mm FWHM). Regional time series were then extracted using AICHA and BNA parcellations. Functional connectivity was computed as the Pearson correlation coefficient between all regional pairs and Fisher z-transformed. Individual connectivity matrices were averaged across subjects to form the group-level normative functional connectome.

The normative functional connectome was used exclusively for CDM analyses as a comparator and was not incorporated into the NDM framework, which relied solely on structural connectomes.

#### Null model implementation

To evaluate whether CDM and NDM results exceeded spatial and topological baselines, three complementary null model classes were implemented.

**Spatial autocorrelation–preserving null model (BrainSMASH)**

Spatially constrained surrogate maps were generated using the BrainSMASH toolbox.[12] For each atlas, parcel-wise Euclidean distance matrices were computed from centroid coordinates. Empirical GMV change maps served as target maps. Surrogate maps were generated using default BrainSMASH parameters (pv = 25, ns = 1), with kNN automatically determined within the neighbor-proportion range ($\delta = 0.1$–$0.9$). A total of 1000 surrogate maps were generated for each analysis.

**Spin-based null model**

Cortical parcels were projected onto an fsaverage spherical surface. Random rotations were applied and parcel values reassigned based on nearest neighbors. Identical rotations were mirrored across hemispheres to preserve symmetry. A total of 1000 rotated surrogate maps were generated using the netneurotools toolbox.[13]

**Topology-preserving network null model**

Topology-preserving surrogate networks were generated using the Maslov–Sneppen rewiring algorithm implemented in the Brain Connectivity Toolbox.[14] This procedure preserves number of nodes, number of edges, and empirical degree sequence

For each surrogate network, five rewiring attempts were performed per edge (bin_swaps = 5), and edge weights were reassigned at frequency 0.1 (wei_freq = 0.1). A total of 1000 surrogate networks were generated.

All three null models (BrainSMASH, spin, and degree-preserving rewiring) were applied in CDM analyses. For NDM analyses, BrainSMASH and degree-preserving rewiring were applied, as NDM simulations operate on structural Laplacians rather than cortical surface embeddings.

#### NDM-Based prediction of GMV atrophy

To determine whether lesion topology alone contains sufficient information to predict individualized diffusion-stage dynamics, ridge regression was used to estimate subject-specific NDM propagation stage (NDM-PS) from lesion-derived regional features.

Ridge regression minimizes the sum of squared prediction errors augmented with an L2 penalty term applied to regression coefficients.[15] This regularization stabilizes solutions in high-dimensional predictor spaces and mitigates multicollinearity. A nested leave-one-out cross-validation (LOOCV) framework was employed.[16] Outer loop: one subject was held out as the test case; the remaining N-1 subjects formed the training set. Inner loop: within each outer training set, an inner LOOCV optimized the regularization parameter λ. Candidate λ values were sampled logarithmically from $2 \times 10^{-10}$ to $2 \times 10^{6}$. For each λ, the model was trained on lesion-derived predictors and empirically derived NDM-PS values, and performance was quantified as the Pearson correlation between predicted and observed NDM-PS across inner folds. The λ yielding the highest correlation was selected for the outer loop. Using the optimal λ, the ridge model was retrained on the full outer training set and used to predict NDM-PS for the held-out subject. This procedure yielded fully out-of-sample predicted NDM-PS values for all subjects.

Overall prediction accuracy was quantified as the Pearson correlation between predicted and empirically derived NDM-PS values across individuals. Statistical significance was assessed via permutation testing. Within each outer fold, NDM-PS values in the training set were randomly permuted 1000 times while keeping lesion predictors fixed. The complete nested cross-validation procedure was repeated for each permutation to generate a null distribution of prediction correlations.

Predicted individual NDM-PS values were subsequently reintroduced into the NDM framework to generate individualized simulated GMV atrophy maps. Spatial correspondence between predicted and observed GMV change maps was quantified using Pearson correlation and compared with correspondence obtained using empirically derived NDM-PS values via paired t-tests.

This framework enables stage-informed prediction of individualized GMV atrophy patterns without reliance on longitudinal follow-up imaging.

### Supplementary results

#### Supplementary Table 1. Demographic and clinical characteristics of participants in the main longitudinal cohort

| | Phase | | |
|---|---|---|---|
| **Characteristics** | **2 weeks** | **3 months** | **12 months** |
| Number of participants | 72 | 69 | 62 |
| Age (years) | 53.93 ± 10.47(22-79) | 53.78 ± 10.65(22-79) | 53.66 ± 10.31(22-77) |
| Time from onset (days) | 13.1 ± 4.8(6-28) | 114.5 ± 20.6(85-187) | 389.7 ± 52.0 (343-749) |
| Sex (male: female) | 36:36 | 35:34 | 27:35 |
| Handedness (right: left) | 66:6 | 63:6 | 57:5 |
| Education (years) | 13.20 ± 2.47(9-20) | 13.14 ± 2.41(9-20) | 13.37 ± 2.52(9-20) |
| Lesion side (right: left) | 30:42 | 29:40 | 26:36 |
| Lesion size ($cm^3$) | 31.32 ± 37.20(0.83-223.21) | 37.71 ± 31.90 (0.83-223.21) | 27.73 ± 29.00(0.83-118.14) |

Values are reported as means ± standard deviations (min–max).

#### Supplementary Table 2. Demographic and clinical characteristics of participants in the independent early-phase cohort

| **Characteristics** | |
|---|---|
| Age (years) | 63.25 ±12.03(37-81) |
| Sex (male: female) | 12:8 |
| Lesion side (right: left) | 8:12 |
| Lesion size ($cm^3$) | 4.10 ± 3.01(0.73-14.09) |
| Two scan intervals (days) | 4 ± 0.5(3-5) |

Values are reported as means ± standard deviations (min–max).

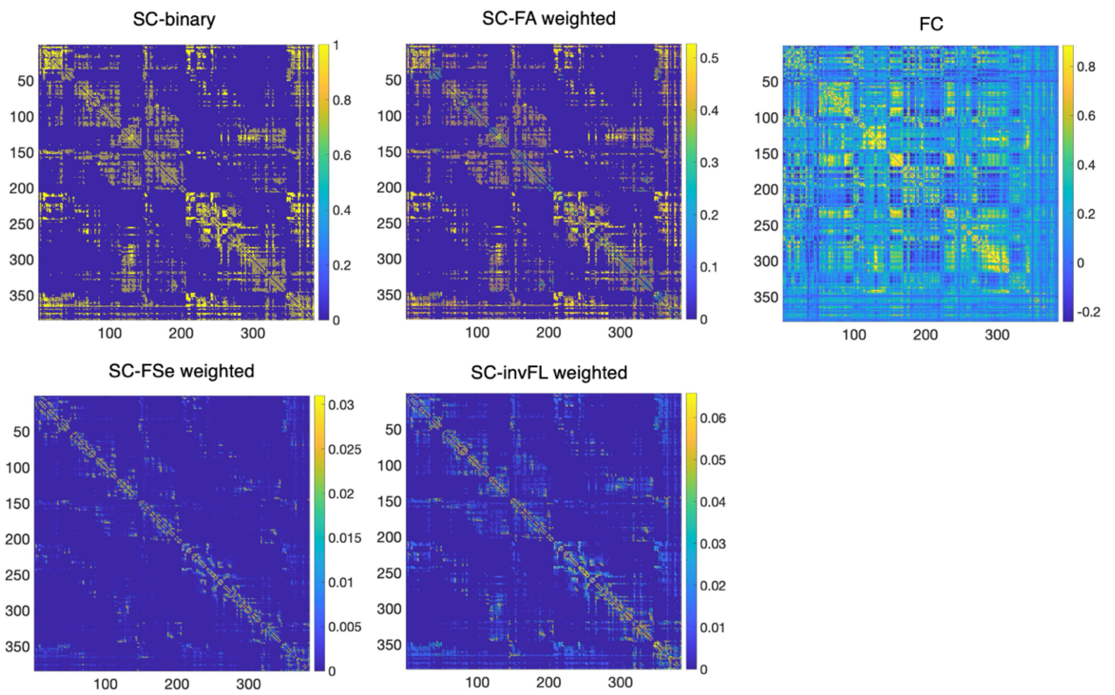


**Supplementary Figure 1. Group-level normative structural and functional connectomes.** Matrices depict the normative connectivity backbone at the AICHA-atlas resolution, derived from healthy cohort data. Color bars indicate the connection strength or probability for each respective weighting metric. SC: structural connectivity; FC: functional connectivity; FA: Fractional Anisotropy; FSe: fraction of streamlines; invFL: inverse fiber length.

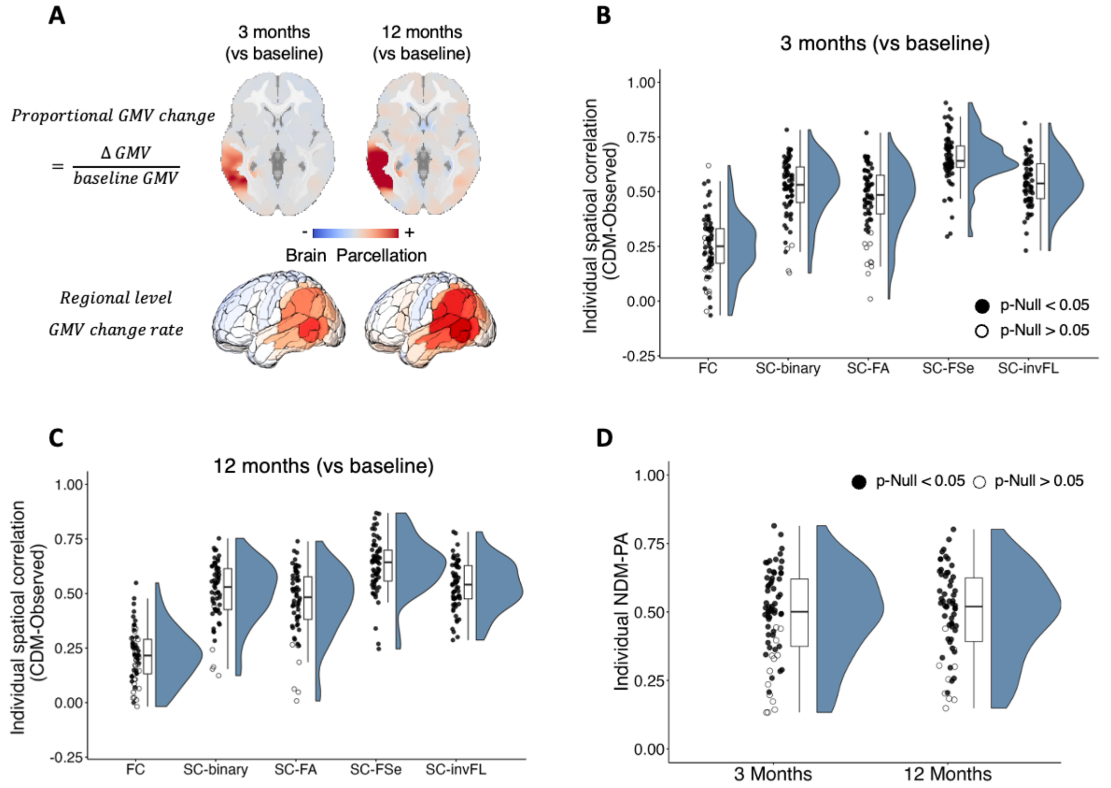


**Supplementary Figure 2. Validation of connectome-constrained and diffusion-based models under BNA atlas.** (A) Spatial maps of proportional GMV change (ΔGMV/ baseline GMV) at 3- and 12-months post-stroke for a representative patient, with regions parcellated via the BNA atlas. GMV decline was most pronounced in peri-lesion regions and extended to remote regions in a spatially structured pattern. (B–C) Spatial correlations between CDM-estimated and observed GMV changes across the entire cohort at (B) 3 months and (C) 12 months post-stroke. Statistical significance was assessed against null models, with filled markers denoting correlations significantly higher than the null distribution ($P_{null} < 0.05$) and open markers indicating non-significant results. Structural connectivity (SC)-based models outperformed functional connectivity (FC)-based models at both time points, with FSe-weighted structural connectivity yielding the strongest correspondence. (D) Distributions of individual maximal NDM–observation correlations. Regional GMV atrophy was simulated using the NDM using the BNA-based structural connectome as the substrate. Spatial correspondence between NDM-predicted and observed atrophy maps was evaluated at 3- and 12-months post-stroke, with filled symbols denoting significant correlations against the null distribution ($P_{null} < 0.05$). NDM captured distributed atrophy patterns at both follow-up time points.

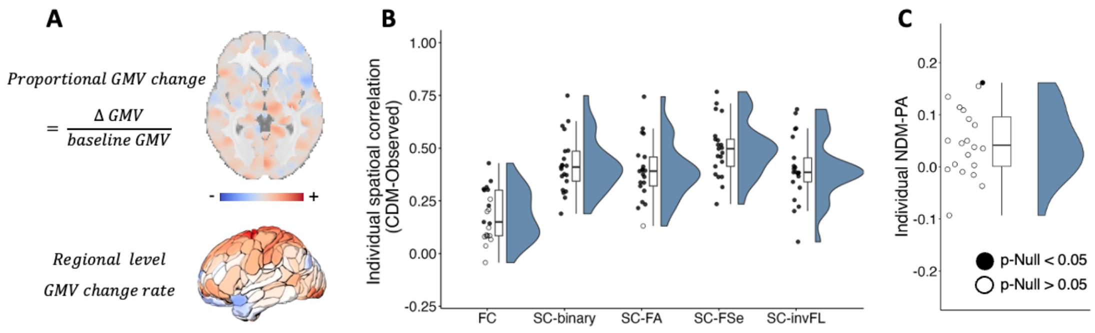


**Supplementary Figure 3. Network-based modeling of GMV atrophy during the hyperacute and early acute stages within the first week after stroke.** (A) Spatial maps of proportional GMV change within the first week. Individualized maps were parcellated using the AICHA atlas. In contrast to chronic stages, GMV alterations within the first week exhibited a relatively uniform distribution, lacking a pronounced lesion-related spatial gradient. (B) Spatial correlations between CDM estimates and observed GMV changes across the cohort. The CDM was applied to evaluate early post-stroke GMV change patterns. While the overall model fit was lower than in long-term transitions, significant variance was captured, with SC consistently outperforming FC. FSe-weighted SC remained the superior predictor, indicating that connectome-constrained redistribution is detectable even at these earliest stages. (C) Distributions of individual NDM–observation correlations within the first week. Regional GMV atrophy was simulated using NDM. For the majority of patients, the correspondence between NDM-simulated and observed patterns did not significantly deviate from null expectations ($P_{null}$ < 0.05). These results suggest that network-mediated atrophy patterns simulated by the NDM become statistically prominent only as the pathological cascade unfolds into subacute and chronic stages.

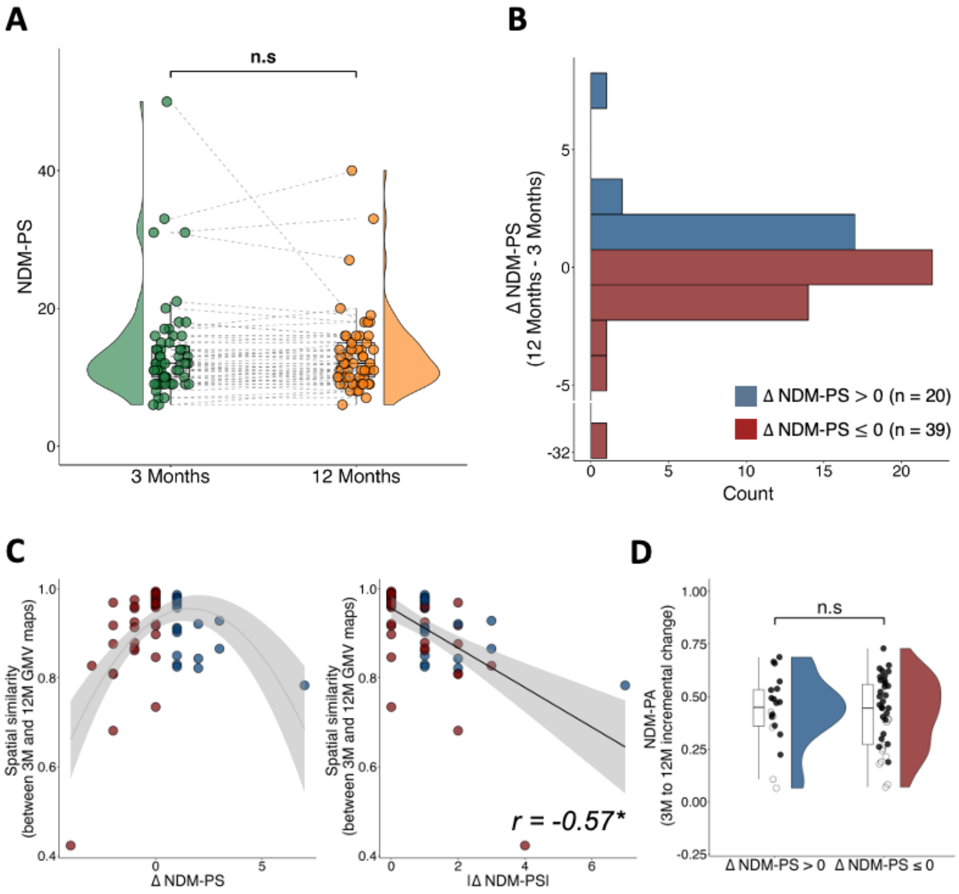


**Supplementary Figure 4. Replication of longitudinal diffusion dynamics under BNA atlas.** (A) Group-level comparison of NDM propagation stage (NDM-PS) derived from BNA-based GMV atrophy patterns at 3- and 12-months post-stroke. Consistent with the main findings, no significant difference was observed between the two time points. (B) Distribution of intra-individual differences in propagation stage ($\Delta$NDM-PS = NDM-PS$_{12\ \text{months}}$ – NDM-PS$_{3\ \text{months}}$). Similar to the primary analysis, $\Delta$NDM are distributed around zero, confirming substantial inter-individual variability in diffusion timing. (C) Spatial similarity between BNA-based 3- and 12-month GMV change maps (left) and its correlation with the absolute difference in stage shifts (|$\Delta$NDM-PS|) across participants (right). (D) Comparison of NDM prediction accuracy (NDM-PA) for GMV changes from 3 to 12 months between the $\Delta$NDM-PS $> 0$ and $\Delta$NDM-PS $\leq 0$ subgroups. Although the trend mirrored the primary results, no statistically significant difference in model correspondence was observed between these two subgroups at this nodal resolution. n.s: non-significant, $^{*}P < 0.05$.

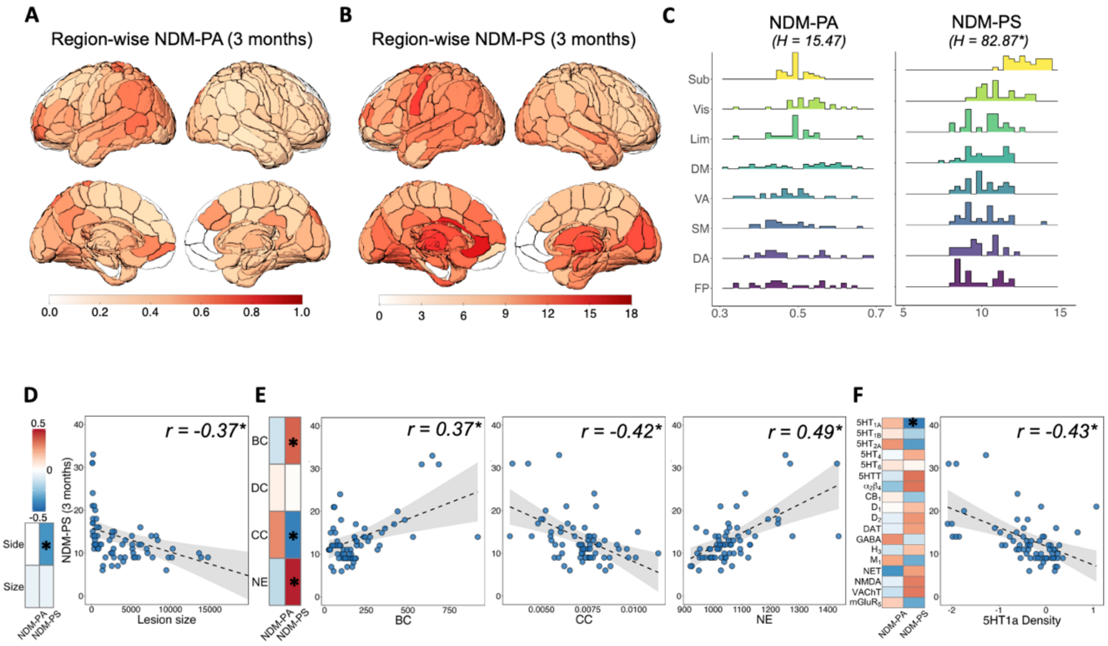


**Supplementary Figure 5. Validation of associations between NDM parameters and lesion-related topology and neurochemical variables using the BNA.** (A-B) Region-wise maps illustrating spatial variation in (A) NDM-PA and (B) NDM-PS as a function of lesion location, calculated using the BNA. Regional values represent the mean NDM metric at 3 months post-stroke across all subjects whose lesions overlapped each specific brain region. (C) System-level distribution of NDM-PA and NDM-PS. Regional NDM metrics are grouped into the seven Yeo functional networks and a subcortical system. Asterisks indicate significant inter-system differences determined by Kruskal–Wallis tests (*P*<0.001), with NDM-PS values being notably higher in the subcortical and visual systems compared to others. Vis, visual; SM, somatomotor; DA, dorsal attention; VA, ventral attention; Lim, limbic; FP, frontoparietal; DM, default mode; Sub, subcortical. (D-F) Subject-level associations of NDM metrics with lesion-related properties. (D) Correlations with global lesion characteristics, confirming the significant association between lesion size and NDM-PS. (E) Correlations with lesion-site network topological properties; besides the consistent correlations with BC and Clu, NE also shows a significant relationship with NDM metrics under the BNA. (BC, betweenness centrality; DC, degree centrality; CC, clustering coefficient; NE, nodal efficiency). (F) Correlations with PET-derived regional densities of 18 neurotransmitter receptors/transporters; notably, the overall correlation patterns remain highly consistent with the results from the AICHA atlas, with 5-HT1a density showing a significant negative association with NDM-PS. In each panel, heatmaps (left) show Pearson correlation coefficients between NDM metrics and the respective features, while scatter plots (right) highlight representative significant correlations. Asterisks denote associations surviving permutation-based FDR correction ($P_{FDR} < 0.05$). Dashed lines and shaded areas represent the linear fit and 95% confidence interval, respectively.

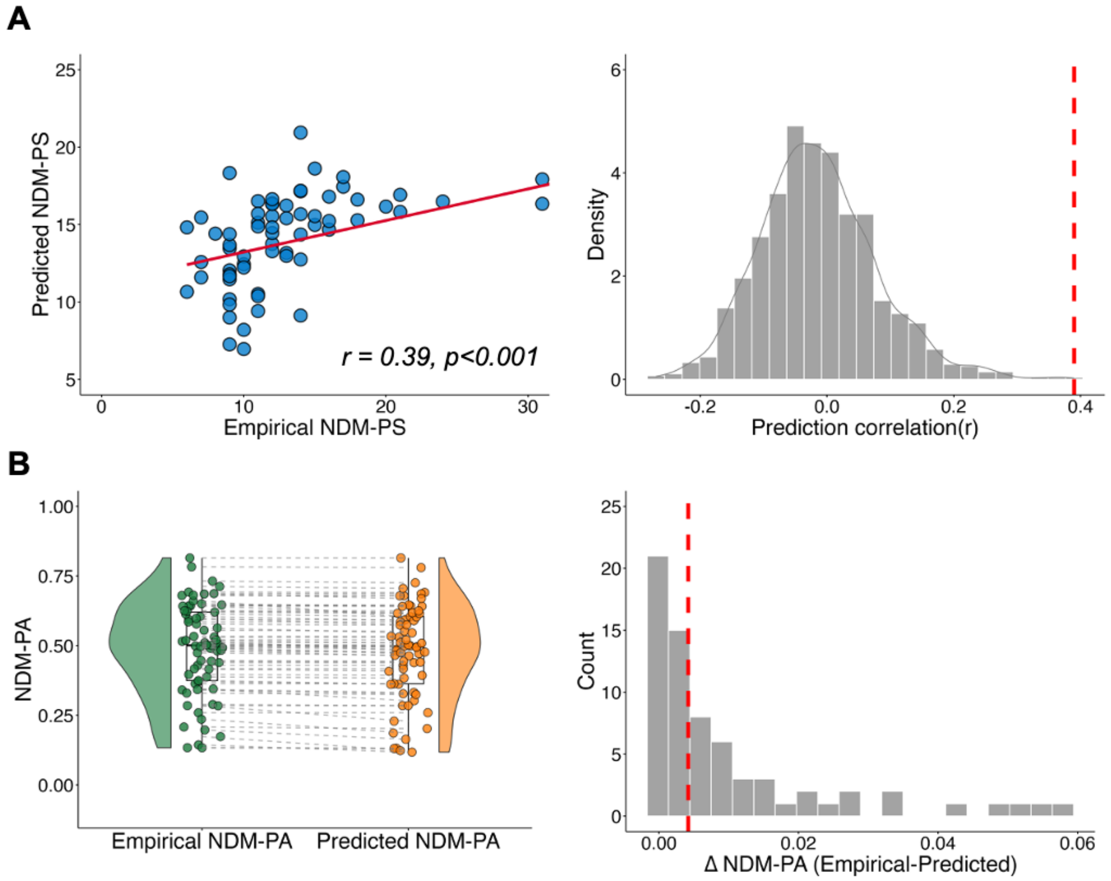


**Supplementary Figure 6. Individual-level prediction of GMV atrophy patterns based on the BNA atlas.** (A) Prediction performance of NDM propagation stage using BNA-based metrics. Left: Pearson correlation between predicted and empirically derived NDM-PS across individuals. Right: null distribution of prediction performance obtained from permutation testing, with the dashed line indicating the observed correlation. The results demonstrate that NDM-PS can be accurately predicted from lesion topography, with the observed correlation significantly exceeding the null model ($p < 0.001$). (B) Spatial correspondence between observed GMV atrophy patterns and NDM predictions generated using empirically derived NDM-PS versus regression-predicted NDM-PS. Left: Group-level distributions of NDM prediction accuracy (NDM-PA) demonstrate comparable spatial performance between the two approaches. Right: the distribution of differences in NDM-PA between empirical-stage and predicted-stage models, with the dashed line indicating the mean difference. Consistent with the main analysis, regression-predicted propagation stage (PS) achieves prediction performance highly similar to that of the empirical model, confirming the robustness of the predictive framework.
